# The interaction between virus-bound KLF4 and host-bound PARP1 directs the localization of Adeno-Associated Virus Type 2 (wtAAV2) to cellular sites of DNA damage

**DOI:** 10.64898/2026.02.24.707757

**Authors:** Rhiannon R. Abrahams, Clairine I.S. Larsen, Raymond Ha, Kinjal Majumder

## Abstract

Wild-type adeno-associated virus type 2 (wtAAV2) is a small, non-pathogenic DNA virus within the family *Parvoviridae* that is modified to engineer recombinant AAV (rAAV) vectors used in gene therapy. wtAAV2 genomes localize to nuclear domains enriched in DNA Damage Response (DDR) factors through mechanisms that remain unknown. We have discovered that the host transcription factor KLF4 (Krüppel-like Factor 4) and the DDR protein PARP1 (poly-ADP ribose polymerase 1) are key effectors regulating wtAAV2 nuclear trafficking. *In-silico* analysis revealed that wtAAV2-associated genomic sites are enriched in KLF4 binding motifs. Confocal imaging demonstrated a close spatial association between KLF4 and wtAAV2 nuclear reservoirs, consistent with KLF4’s known ability to form nuclear condensates. Mutation of the KLF4 binding site within the wtAAV2 genome and RNAi-mediated knockdown of KLF4 globally reduced the expression of viral *Rep68/78* genes and attenuated the localization of the viral genome to cellular DDR sites. Chemical inhibition of the KLF4-interacting protein PARP1 with Olaparib decreased the ability of wtAAV2 genomes to localize to cellular DDR sites and transcribe viral genes. Ectopic expression of wild-type PARP1, but not its KLF4-binding-deficient mutant, rescued wtAAV2 gene expression in PARP1-deficient cells. These findings define a novel mechanism by which wtAAV2 exploits the interaction between host transcription factors and DNA repair machinery to establish a persistent nuclear niche. Insertion of KLF4-binding elements into recombinant AAV2 (rAAV2) gene therapy vectors is sufficient to enhance transduction of target cells, providing a framework for engineering vectors with improved nuclear targeting and transcriptional activity.

**IMPORTANCE:** Wild-type Adeno-associated virus type 2 (wtAAV2) has emerged as the preferred platform for engineering gene therapy vectors due to its non-pathogenic nature and ability to persist in host cells long-term. However, limited understanding of how wtAAV2 genomes navigate the nuclear environment to establish viral reservoirs has hindered the development of efficient recombinant AAV (rAAV) vectors. We demonstrate that the protein KLF4 bound to wtAAV2 genome recruits the virus to cellular KLF4 sites bound by PARP1. Disruption of either KLF4 binding, PARP1 activity or KLF4-PARP1 interaction significantly impairs wtAAV2 localization and transcription, highlighting the importance of their function in the non-replicative wtAAV2 life cycle. KLF4 binding sites are sufficient to improve the expression of transgenes from rAAV vectors and increase their association with cellular DDR proteins. This study advances our understanding of wtAAV2-host interactions and opens new avenues for improving rAAV gene therapy platforms.

## INTRODUCTION

DNA viruses have evolved complex strategies to express viral genes, amplify their genomes, persist within the host-cell nucleus and induce cell death. To accomplish this, they usurp cellular transcription, replication and DNA processing factors while evading elimination by cellular defense pathways^1, 2^. Many DNA viruses exploit the host DNA Damage Response (DDR) pathways, a redundant network of signaling proteins that are tasked with sensing and processing DNA lesions to maintain the fidelity of the genetic code^3–5^. These interactions allow DNA viruses to remodel nuclear subdomains, redirect repair machinery, and create microenvironments that support viral persistence or replication. By co-opting DDR proteins, viral genomes establish favorable niches that support viral life cycle, be it for replication [as is the case for lytic DNA viruses like Minute Virus of Mice (MVM)^6, 7^ and SV40^8–11^], or long term persistence [as is the case for the latent phase of DNA viruses like Epstein-Barr Virus (EBV)^12–14^ or Herpes Simplex Virus (HSV)^15^]. The genomes of replicating and non-replicating wild-type Adeno-Associated Virus Type 2 (wtAAV2) associate with cellular sites of DNA damage using mechanisms that remain unknown^16^. Interestingly, recombinant AAV2 (rAAV2) genomes do not associate with cellular DDR sites efficiently, suggesting these interactions are a key part of the viral life cycle^16, 17^.

The cause-effect relationship between DNA viruses and cellular DDR pathways represents an exciting frontier in understanding virus-host genome interactions. Tumor viruses like human papillomavirus genomes are anchored to cellular DDR sites using the host transcription factor BRD4 and viral E2 proteins^18–20^. Several of these genomic regions are super-enhancer elements which are prone to undergo DNA breaks due to chromatin accessibility and high local transcriptional activity^21, 22^. Genomes of Hepatitis B Virus (HBV) are also associated with cellular DDR sites, many of which are promoter-proximal and transcriptionally active^23^. These observations suggest a shared relationship in which viral genomes preferentially localize to host chromatin regions where transcription, chromatin landscape, and DNA repair pathways intersect. The autonomous parvovirus MVM localizes to cellular DDR sites (which also correlate with transcriptionally active Type A chromatin) using viral NS1 bound to the viral genome^7, 24^. Interestingly, NS1’s ability to localize to cellular DDR sites utilizes the cellular DDR signaling kinase ATR^25^. However, the mechanism of wtAAV2 genome localization during non-replicative infection remains unknown.

Infection of host cells with wtAAV2 induces cellular replication stress via RPA exhaustion, likely through molecular mimicry, where the wtAAV2 genome resembles a stalled replication fork^17, 26^. The formation of single-stranded DNA at stalled host replication forks is recognized by cellular Poly-ADP-Ribose Polymerase 1 (PARP1), leading to its activation^27^. The ensuing cellular DNA damage might create fertile regions such as fragile sites for parvovirus genomes to localize^28, 29^. PARylation, which adds Poly-ADP-ribose polymers to generate a scaffold around the DNA break site, plays a multifaceted role in the nuclear environment that can potentially be usurped by DNA viruses, including ITR-containing rAAV2 genomes^30^. In support of this model, PARylation of host architectural proteins like CTCF leads to the maintenance of open chromatin on EBV genomes^31–33^. Cellular PARylation levels can be associated with DDR pathway proteins like DNA-PKcs and MRE11, which are also known to regulate wtAAV2-induced signals^34, 35^. In response to genotoxic stress, PARP1 modifies the cellular transcription factor KLF4 (Kruppel-Like Factor 4), facilitating its association with chromatin to regulate transcription^36, 37^. DNA viruses that activate cellular DDR signals like HPV and EBV use KLF4’s role in differentiation-dependent amplification of host cells to also amplify the viral genome^38, 39^. However, whether KLF4 plays a role in the nuclear persistence of the non-replicative wtAAV2 genome remains unknown.

The wtAAV2 non-structural proteins REP68 and REP78, which are generated at low levels during wtAAV2 mono-infection, are known to associate with PARP1 without the necessity of a DNA intermediate^40^. PARP1 associates with wtAAV2 and rAAV2 genomes at their ITR elements, rendering the host genome vulnerable to replication stress and subsequent DNA damage that can lead to cell death^30^. Proteomics analysis of the eukaryotic replisomes in wtAAV2 infected cells have previously revealed a depletion of cellular PARG, which is tasked with countering the effects of PARylation on the host genome^17, 41, 42^. These observations indicate that the interactions between wtAAV2 and PARP1 at cellular DDR sites might play a key role in determining the outcomes of viral infection and long-term success of rAAV2 gene therapy vectors. Addressing these gaps in knowledge of how wtAAV2 genomes are targeted to DDR-rich regions, how viral genomes persist long-term at distinct nuclear sites, and how the cellular DDR factors are usurped to process virus genomes are critical for improving rAAV vector effectiveness.

In this study, we investigated how wtAAV2 genomes navigate the nuclear environment and exploit cellular DDR factors to establish viral reservoirs at distinct genomic sites. wtAAV2 genomes associate with cellular DDR sites that are enriched in binding elements for cellular KLF4. The wtAAV2 genomes also contain KLF4 binding sites upstream of all three viral promoters P5, P19 and P40. Global knockdown, as well as deletion of the KLF4 binding site on wtAAV2 leads to decreased viral gene expression and attenuated localization of the viral genome to cellular DDR sites. This coincides with decreased PARP1 association with the wtAAV2 genome. Interestingly, insertion of the KLF4 binding element into rAAV2 vectors is sufficient to enhance transgene expression and association with cellular DDR factors. Taken together, these findings uncover a novel interaction between cellular DDR proteins and key transcription factors that are usurped by wtAAV2 to regulate the fate of virus genomes in the nucleus.

## RESULTS

### KLF4 binding elements are enriched at wtAAV2-associated cellular DDR sites and on wtAAV2 genome

The spatial interaction between parvovirus genomes and cellular DDR sites seems to play a key role in regulating the viral life cycle. We have previously observed that replicating and non-replicating wtAAV2 genomes associated with cellular sites of DNA damage^16^ marked by phosphorylated histone H2AX [referred to as γH2AX^43^]. To validate that nonproductive wtAAV2 genomes localize to induced sites of DNA damage, we performed laser micro-irradiation assays in U2OS cells during wtAAV2 infection at 24 hours post-infection (hpi) by irradiating focused subnuclear regions. We monitored the relocalization of the wtAAV2 genomes using immunofluorescence imaging coupled with fluorescent in-situ hybridization (Immuno-FISH). In these studies, induced DNA break sites were immunolabelled with a γH2AX antibody following hybridization of fluorophore-conjugated primers (Table 1) complementary to the wtAAV2 genome. As shown in the representative Immuno-FISH images of a single cell, the γH2AX (red) and wtAAV2 genomes (green) formed punctate foci in un-irradiated cells (Fig. 1A, top). These fluorescent labels rapidly relocalized to induced DNA breaks observable as a stripe (Fig. 1A, bottom). Quantification of wtAAV2 signal intensity shows colocalization of the viral genomes with induced DNA damage sites across the stripe (Fig. 1B).

**Figure 1.**
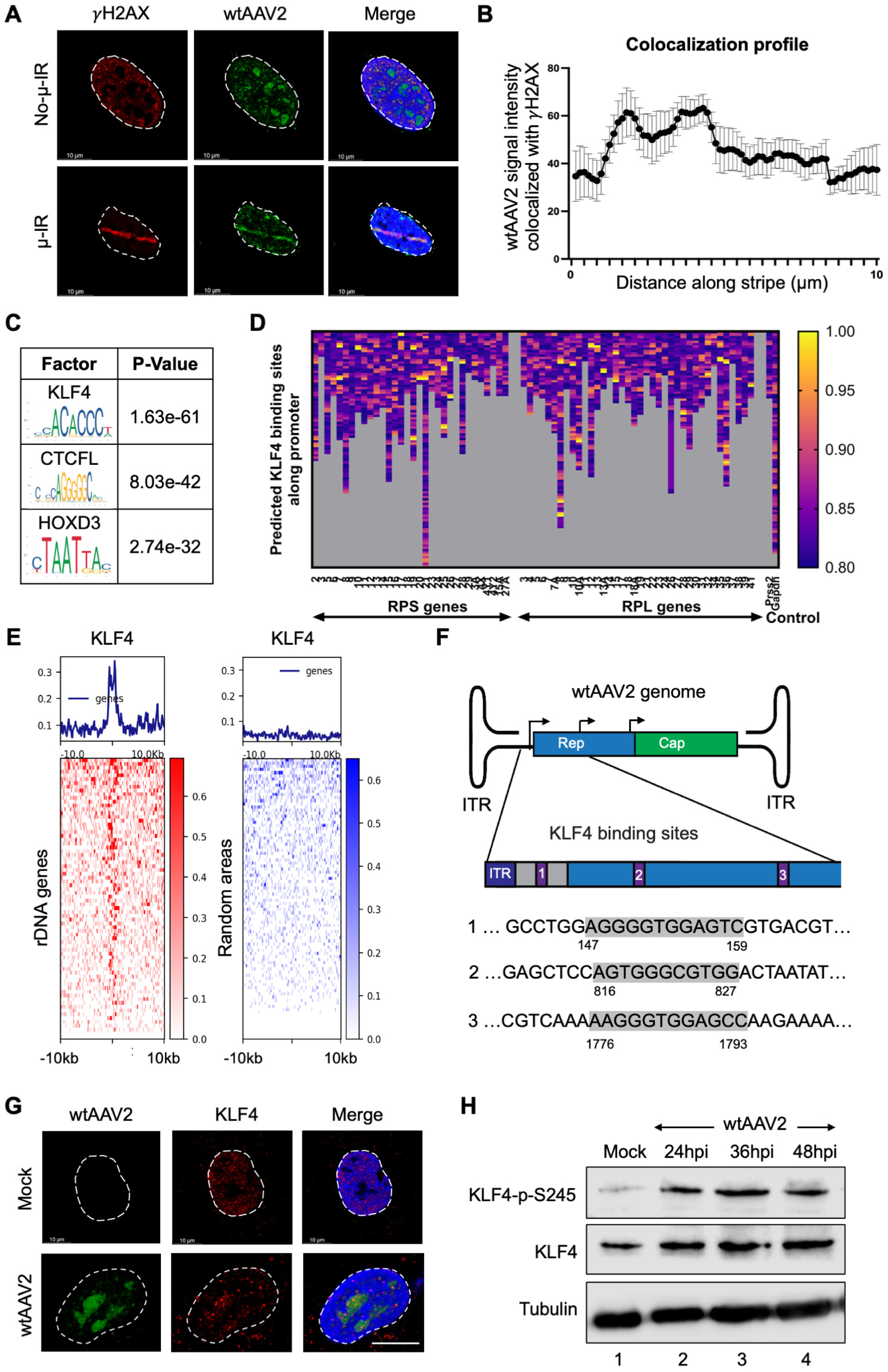
KLF4 binding elements are enriched at wtAAV2-associated cellular DDR sites and on wtAAV2 genome. (A) Immuno-FISH analysis of wtAAV2-infected U2OS cells at an MOI of 5,000 vg/cell at 24 hpi where cellular DNA damage was induced by laser micro-irradiation. The location of the wtAAV2 genome was monitored by FISH using primers complementary to the wtAAV2 genome (green) and the DNA damage site was assessed using γH2AX staining (red). DAPI represents the nucleus (blue) with white dashed line demarcating the nuclear border. The white scale bar represents 10 microns. (B) Average of wtAAV2 signal intensity that colocalized with the γH2AX staining in multiple nuclei averaged over 10 microns of laser striping divided into 75 smaller bins. Data is presented as mean ± SEM of at least 10 independent laser micro-irradiated cells. (C) Results of the *in-silico* prediction analysis of host transcription factors binding sites that were enriched at wtAAV2-associated genomic region (computed using JASPAR informatics resource^83^). The most common consensus motifs (leftmost column) were compared with the p-value of their enrichment in the given regions (rightmost column). (D) Outcomes of *in-silico* analysis of KLF4 binding sites in the promoter regions of ribosomal genes that make up the nucleolus. The columns represent the number of KLF4 binding elements detected in each of the indicated gene promoters with the heatmap intensities depicting the strength of predicted KLF4 binding. (E) Analysis of the presence of KLF4 binding sites (assessed by KLF4 ChIP-seq in MCF7 cells) within the 10 kb windows spanning the promoters of ribosomal DNA genes (left; red heatmaps) compared with control sites on the host genome generated randomly (right; blue heatmaps). The profiles of the binding elements are shown on the line graph above each respective heatmap and was computed using deeptools on the Galaxy platform^84^. (F) Schematic of the wtAAV2 genome with the promoters and respective KLF4 binding sites demarcated in the inset. The zoomed-in insets indicate the KLF4 binding sequences highlighted in grey and their positions on the wtAAV2 genome. (G) Immuno-FISH images of U2OS cells infected with wtAAV2 at an MOI of 5,000 vg/cell at 24 hpi showing the relative location of KLF4 (red) with that of the viral genome (green). The nucleus is demarcated by DAPI (blue stain) and the nuclear borders are marked by dashed white lines. The scale bar represents 10 microns. (H) Western blot analysis of the impact of wtAAV2 infection of 293T cells at the indicated timepoints on the global KLF4 levels (middle) and phosphorylated KLF4 at serine 245 (top blot). Tubulin levels serve as loading controls for the immunoblots.

**Table 1:**
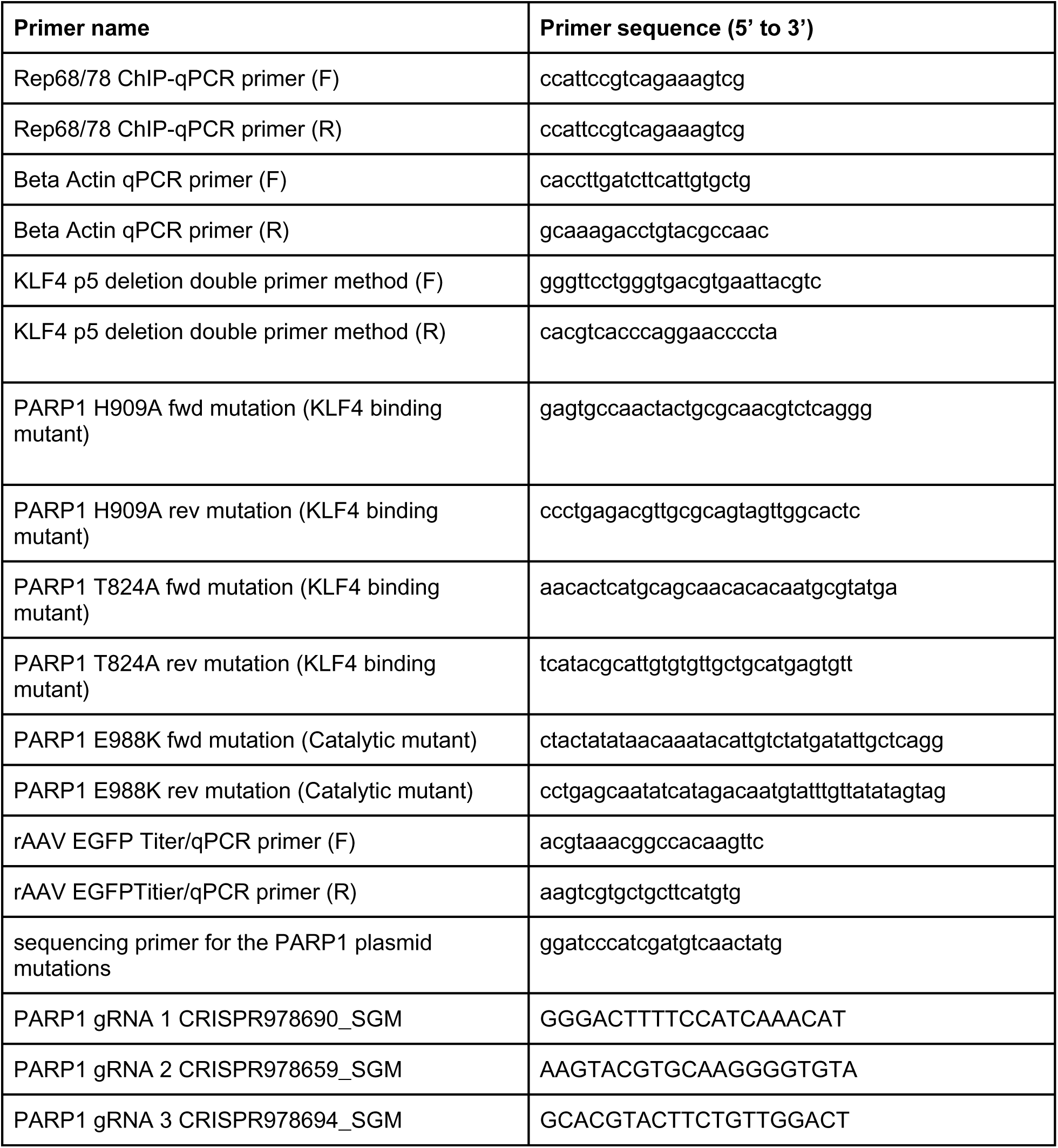
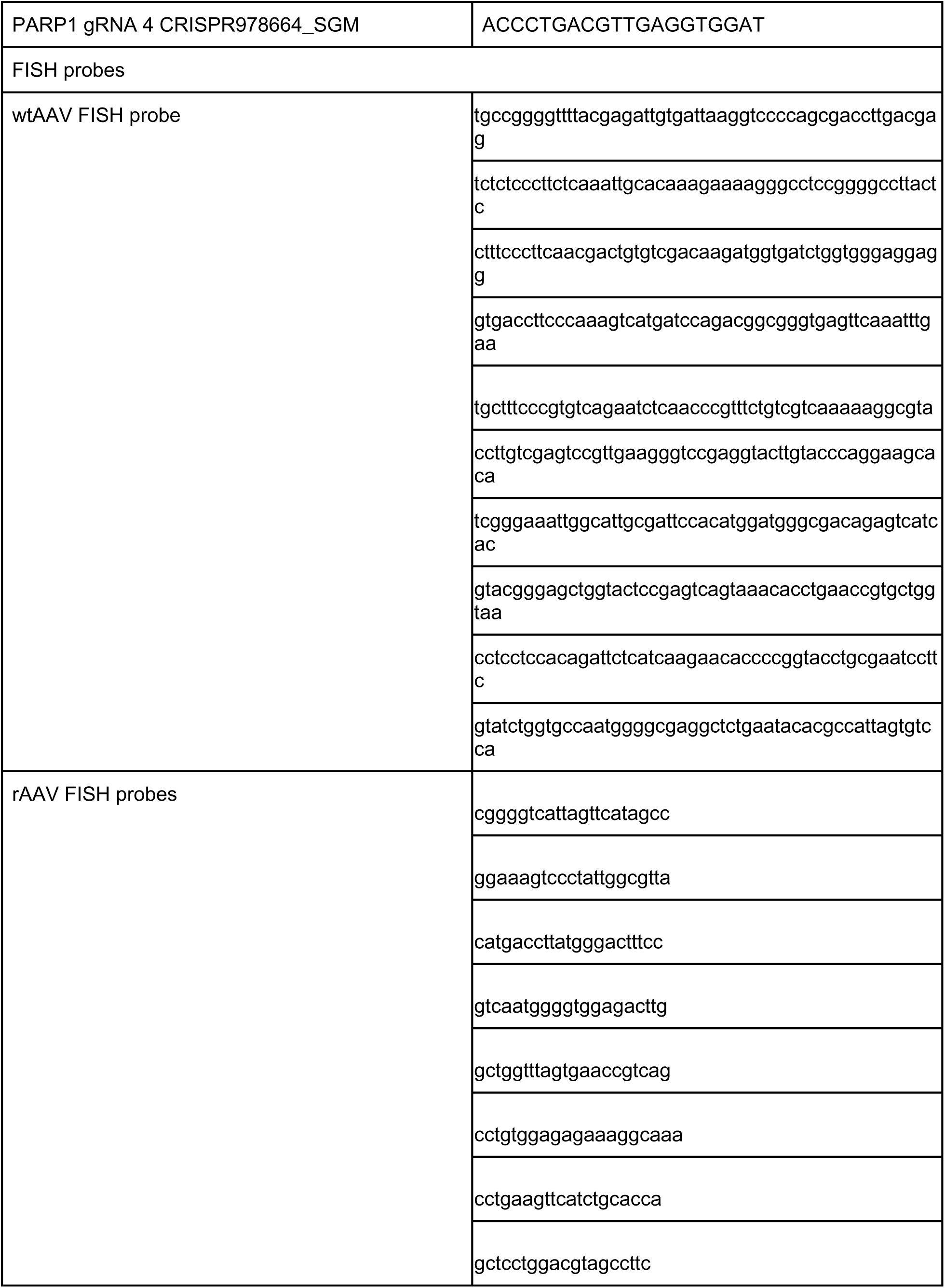
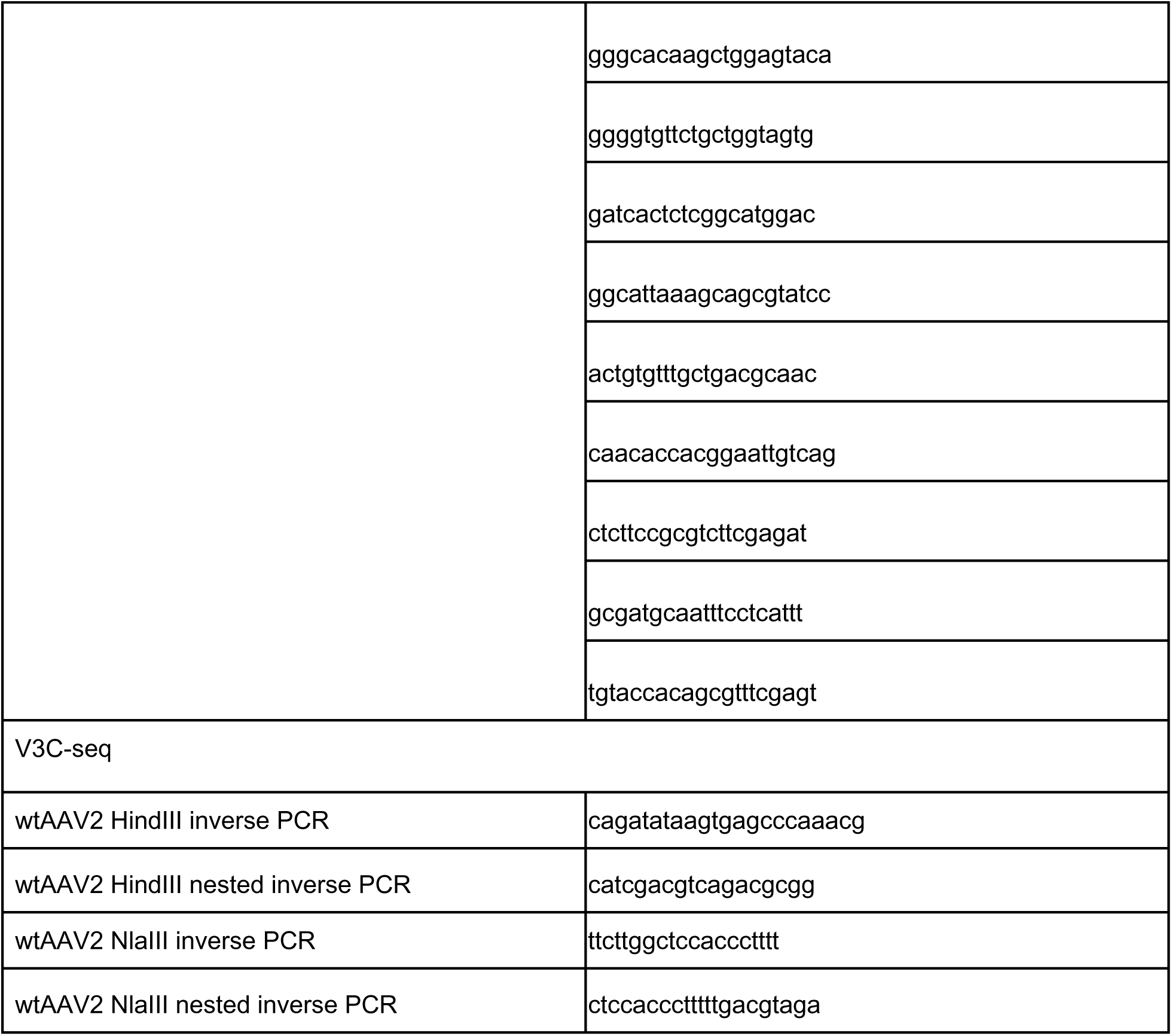
Table of primer and guide RNA sequences.

High-throughput monitoring of the location of the wtAAV2 genome showed that viral genomes were associated with cellular γH2AX sites as distinct genomic peaks instead of broad chromatin domains, which are the characteristic profile of γH2AX-marked genomic DNA (previously published by us^16^ and represented in Fig. 4A). This pattern suggested that wtAAV2 genomes target certain sites of DNA damage on the host. To investigate which host transcription factor binding motifs might be enriched at wtAAV2-associated cellular sites, we selected the fifty genomic regions most strongly associated with wtAAV2 at 24 hours post-infection (hpi) in 293T cells and performed *in-silico* analysis using the MEME suite^44^. As shown in Fig. 1C, motif searching revealed that binding motifs associated with transcription factors KLF4 (p-value 1.63e-61), HOXD3 (p-value 2.74e-32), and architectural protein CTCFL (p-value 8.03e-42) were significantly enriched at wtAAV2-associated cellular genomic sites. Since wtAAV2 genomes have also been shown to colocalize with nucleoli^44^, which are sites of cellular ribosomal gene expression^45^, we examined the enrichment of KLF4 binding elements in the promoter regions of rDNA genes. As shown in Fig. 1D, promoters of rDNA genes on the human genome contain strong KLF4 binding sequences (indicated by white boxes in the heatmaps representing rDNA promoter elements), whereas control genes such as *Gapdh* and *Prss2* lacked strong KLF4 binding sites. Experimental assessment of KLF4 binding sites on rDNA genes in published ChIP-seq datasets^46^ supported our *in-silico* analysis, revealing enrichment of KLF4 binding in the 10 kilobase windows on either side of rDNA promoters (Fig. 1E, left). However, non-rDNA regions of the human genome did not show an enrichment within these 20 kilobase bins (Fig. 1E, right). These findings indicated that wtAAV2-associated cellular loci are enriched for KLF4 occupancy (in addition to having DDR markers).

*In-silico* prediction analysis using the JASPAR platform additionally revealed that KLF4 binding motifs are present upstream of all three wtAAV2 promoters, P5, P19 and P40 (Fig. 1F). KLF4’s ability to form nuclear condensates that bring together distal genomic regions^47^ into the same milieu made it an attractive target that might coalesce KLF4-bound wtAAV2 with KLF4-bound cellular DDR sites. To experimentally test whether wtAAV2 genomes and KLF4 proteins occupy the same nuclear territories, we performed Immuno-FISH studies in wtAAV2 infected U2OS cells at 24 hpi. In these experiments, KLF4 proteins were immunolabelled prior to hybridization of fluorophore-conjugated primers complementary to the wtAAV2 genome. As shown in Fig. 1G, KLF4 proteins were in the vicinity of wtAAV2 genomes in U2OS cells and many of these KLF4 foci colocalized with wtAAV2. Furthermore, wtAAV2 infection time-courses in 293T cells led to the phosphorylation of KLF4 at Serine 245 (KLF4-S245; Fig. 1H), which is activated by cellular ATM/ATR signaling kinases^38, 39^. Taken together, these observations indicated that there is a close spatial association between cellular KLF4 and wtAAV2 genomes that warrants further investigation.

### Global KLF4 knockdown attenuates wtAAV2 localization to DDR sites and viral gene expression

To determine whether KLF4 regulates wtAAV2 life cycle, we targeted KLF4 transcripts using RNAi knockdown in 293T cells and U2OS cells for 24 hours prior to infection with wtAAV2 at an MOI of 5,000 vg/cell (representative knockdown shown in Fig. 2A, left). The intensity of KLF4 relative to Tubulin levels in three independent replicates indicated a significant decrease of KLF4 protein in these cells in response to these siRNAs (Fig. 2A, right). Reverse transcription coupled with quantitative PCR (RT-qPCR) monitoring viral transcript levels revealed at least a 3-fold decrease in *Rep68/78* levels in both cell types transfected with the KLF4-targeting siRNA (Fig. 2B). These observations suggested that KLF4 is required for optimal wtAAV2 gene expression. To determine whether KLF4 regulates the nuclear localization of wtAAV2 genomes to induced DDR sites, we performed laser micro-irradiation assays in U2OS cells transfected with the KLF4 siRNAs before being infected with wtAAV2 as described above. At 24 hpi, distinct regions of the nucleus were irradiated with a confocal laser and immediately processed for imaging to determine the location of the wtAAV2 genome relative to the DDR marker monitored by PARylation staining. As shown in the representative images in Fig. 2C (top panels) and quantified as signal intensity colocalization to PARylated substrates in Fig. 2D, induction of cellular DNA damage by laser micro-irradiation led to relocalization of the wtAAV2 genome to the cellular DNA break sites. However, in KLF4-depleted cells, wtAAV2 re-localization to these induced break sites was severely attenuated. wtAAV2 genomes in these cells instead persisted as nuclear foci in subnuclear spaces distant from the laser-micro irradiated stripes (Fig. 2C, middle and bottom panels). Moreover, these observations were phenocopied in multiple independently irradiated cells when quantified (Fig. 2D). Together, these observations suggested that, on a global scale, KLF4 assists in regulating the localization of wtAAV2 genomes to cellular DDR sites and promotes efficient viral gene expression.

**Figure 2.**
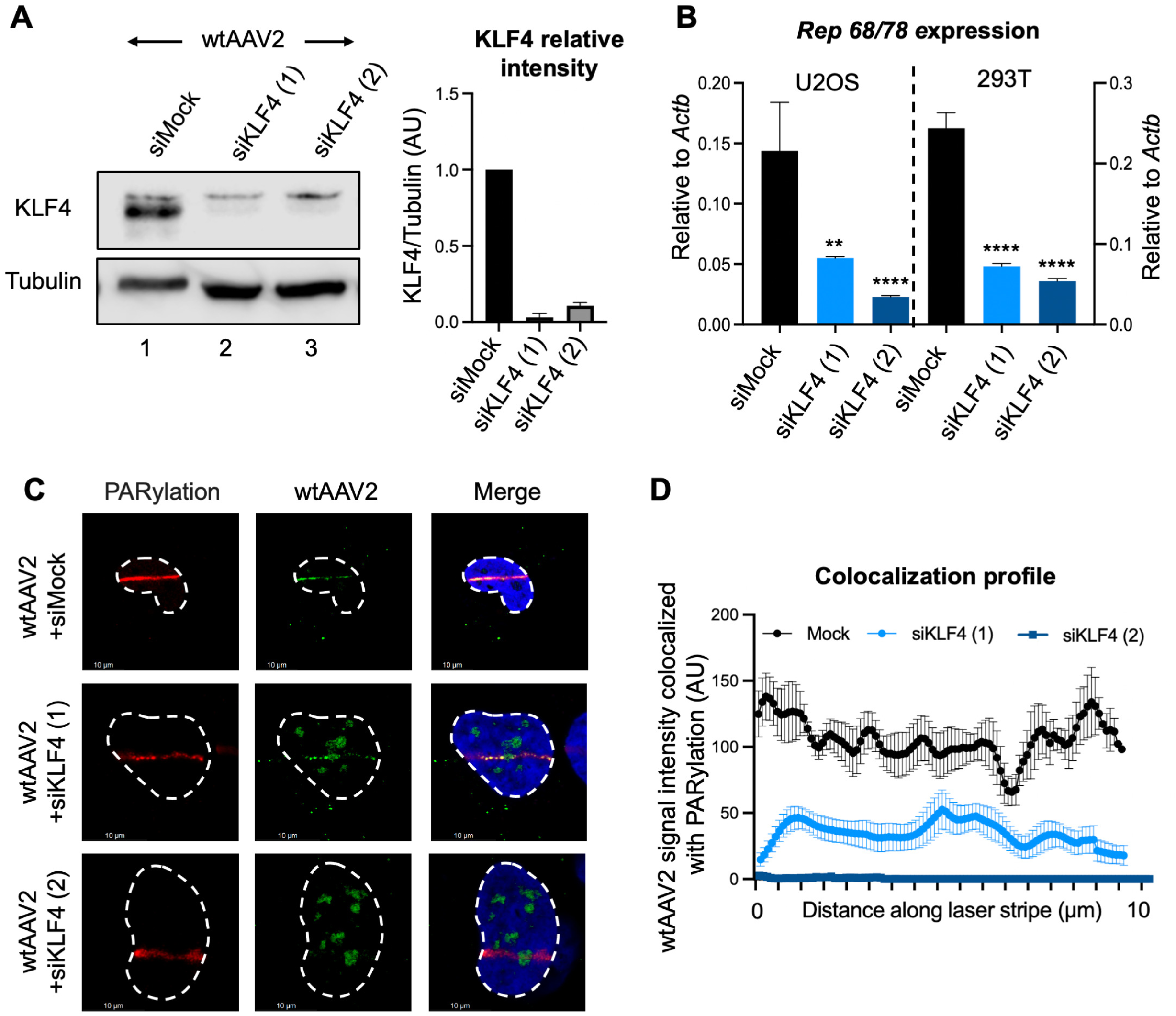
Global KLF4 knockdown attenuates wtAAV2 localization to DDR sites and viral gene expression. (A) Western blot analysis confirming KLF4 knockdown in U2OS cells transfected with two independent siRNAs targeting KLF4 mRNAs compared with siMock–transfected cells infected with wtAAV2 at an MOI of 5,000 vg/cell for 24 hours. Total cellular Tubulin levels were used as loading control. The right half represents quantification and the mean ± SEM of the KLF4 levels relative to Tubulin in three independent replicates. (B) Quantification of *Rep68/78* transcripts relative to *Actb* transcripts in wtAAV2-infected U2OS cells (left) and 293T cells (right) at 24 hpi after knockdown for 24 hours. Data represents the mean ± SEM of three independent experiments with the statistical significance computed using t tests, ** representing p < 0.01, **** representing p < 0.0001. (C) Representative images of the impact of KLF4 knockdown on wtAAV2 localization to induced cellular sites of DNA damage evaluated using laser micro-irradiation coupled with Immuno-FISH in wtAAV2 infected U2OS cells at 24 hpi. The DNA damage sites were monitored by PARylation staining (red) and the viral genomes were tracked by Alexa-Fluor-488-labelled oligos complementary to the wtAAV2 genome (green). The nucleus was monitored by DAPI staining (blue) and the nuclear borders were demarcated by dashed white lines. The white scale bar represents 10 microns. (D) The profile of the intensity of wtAAV2 signal (green) that colocalized with DNA damage monitored by PARylation signal (red) in Immuno-FISH assays across 20-30 nuclei and averaged over the length of the laser micro-irradiated stripe by dividing into 90 bins. Data represents the mean ± SEM of the colocalized intensity at the computed site along the micro-irradiated stripe.

### KLF4 binding site deletion decreases wtAAV2 gene expression and localization to induced cellular DDR sites

To determine whether KLF4 binding to the wtAAV2 genome directly regulates viral life cycle, we generated a mutant clone of wtAAV2 where the KLF4 binding element upstream of the P5 promoter was deleted by site-directed mutagenesis (Fig. 3A, hereafter labelled as wtAAV2^ΔKLF4^). Full plasmid sequencing confirmed that only the KLF4 binding site was deleted in the pAAV2^ΔKLF4^ infectious clone used to generate the mutant virus. Virus was generated by transfection of the wtAAV2 or the wtAAV2^ΔKLF4^ plasmids in producer 293T cells in the presence of pHelper plasmid [expressing Adenovirus E2A, E4 and VA-RNA^48–50^]. Viral titer measurements showed that both infectious clones produced virus particles at equal levels (Fig. 3B). ChIP-qPCR analysis on the viral genome demonstrated a 3-fold reduction of KLF4 binding to the wtAAV2^ΔKLF4^ genome (Fig. 3C) and a 5-fold reduction of PARP1 binding (Fig. 3D). The KLF4 binding sites retained on the P19 and P40 promoter regions might account for the residual KLF4 detected on the wtAAV2^ΔKLF4^ genome. Infection of U2OS and 293T cells with wtAAV2^ΔKLF4^ at an MOI of 5,000 vg/cell at 24 hours revealed a 7-fold decrease of *Rep68/78* transcripts (Fig. 3E). These findings suggested that loss of KLF4 binding regulates wtAAV2 gene expression.

**Figure 3.**
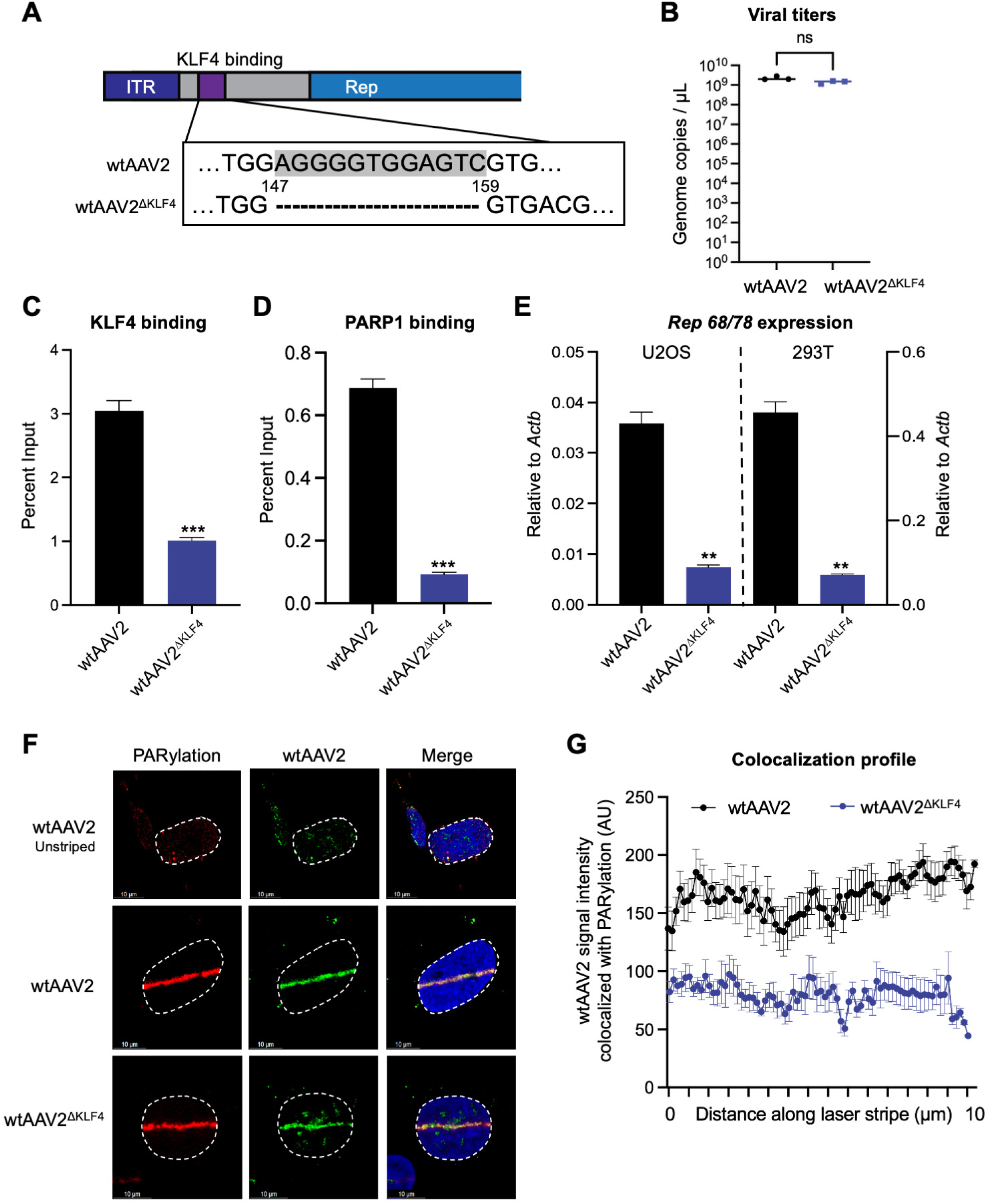
KLF4 binding site deletion decreases wtAAV2 gene expression and localization to induced cellular DDR sites. (A) Schematic of the wtAAV2 genome with the indicated KLF4 site mutation that was generated by site-directed mutagenesis. The grey highlighted sequence in the wtAAV2 region indicates the KLF4 binding sequence. (B) The impact of KLF4 binding element mutation on the viral genome titers produced by 293T cells was monitored using qPCR in three independent replicates. (C) The impact on KLF4 binding and (D) PARP1 binding on the wtAAV2^ΔKLF4^ genome was measured using ChIP-qPCR with primers complementary to the P5 region of the viral genome. ChIP-qPCR data is presented as mean ± SEM of percent input from 3 independent replicates of 293T cell infection with statistical significance computed using t tests. P values represent statistical significance, with *** representing p < 0.001. (E) The impact of KLF4 binding site deletion on *Rep68/78* gene expression was measured using RT-qPCR in U2OS cells (left) and 293T cells (right) at 24 hpi after infection with an MOI of 5,000 vg/cell. Data represents the mean ± SEM of three independent experiments with the statistical significance relative to wtAAV2 computed using t tests, ** representing p < 0.01 and **** representing p < 0.0001. (F) Impact of KLF4 site deletion on wtAAV2 localization to the cellular sites of DNA damage was evaluated using laser micro-irradiation coupled with Immuno-FISH in wtAAV2 infected U2OS cells at 24 hpi. The DNA damage site was monitored by PARylation staining (red) and the viral genomes were monitored by fluorophore-labelled oligos complementary to wtAAV2 (green). The nucleus was monitored by DAPI staining (blue) and the nuclear borders are demarcated by dashed white lines. The white scale bar represents 10 microns. (G) Profile of the intensity of wtAAV2 signal (green) that colocalizes with DNA damage monitored by PARylation signal (red) in Immuno-FISH assays across at least 10 nuclei and averaged over the length of the laser micro-irradiated stripe. Data represents the mean ± SEM of the colocalized intensity at the computed site along the micro-irradiated stripe divided into 75 bins.

To assess how KLF4 binding regulates the relationship between cellular DDR induction and wtAAV2 localization, we performed laser micro-irradiation followed by Immuno-FISH in wtAAV2-infected U2OS cells at 24 hpi. As shown in Fig. 3F and quantified in Fig. 3G, the relocalization of wtAAV2^ΔKLF4^ to induced cellular DDR sites (monitored by PARylation, red) was significantly diminished in wtAAV2^ΔKLF4^-infected cells compared to wtAAV2-infected cells. However, this decrease in relocalization was not completely lost. This partial phenotype might be due to the continued presence of KLF4 binding sites on wtAAV2^ΔKLF4^ genomes (described above). Taken together, our findings suggested that the ability of KLF4 binding elements to interact with the wtAAV2 genome regulates their association with cellular DDR sites and efficient viral gene expression.

### KLF4 binding site deletion attenuates wtAAV2 localization to DDR sites genome-wide

To determine how KLF4 binding site mutation impacts wtAAV2’s localization to cellular DDR sites, we performed a high-throughput chromosome conformation capture assay coupled with sequencing with the viral genome as the viewpoint (location of the inverse PCR primers shown in Fig. 4A, top). The viral chromosome conformation capture coupled with sequencing uses the principle of formaldehyde-based crosslinking to freeze the interaction of wtAAV2 genomes to spatially proximal cellular genomic regions. Using subsequent restriction enzyme digest followed by intramolecular hybrid junction formation, these assays generate novel hybrid DNA molecules between the virus and host genomes. Subsequent high-throughput sequencing identifies the cellular sites proximal to the virus and the frequency with which the virus associates with these genomic regions. We have previously profiled the localization sites of wtAAV2 to the host in 293T and U2OS cells^16^. Comparison of wtAAV2 localization to host genomic sites with that of wtAAV2^ΔKLF4^ virus revealed distinct genomic regions that were shared between wtAAV2 and wtAAV2^ΔKLF4^ in the vicinity of KLF4 binding elements and γH2AX-associated sites (example in Fig. 4A, middle). In contrast, wtAAV2^ΔKLF4^ localization was lost to other host genomic sites (example in Fig. 4A, bottom). Genome-wide comparison of wtAAV2 and wtAAV2^ΔKLF4^ localization (illustrated in the ideograms in Fig. 4B) revealed that 30% of the top 150 wtAAV2-associated sites were shared with the top 150 wtAAV2^ΔKLF4^-associated sites (Fig. 4C). This coincidence of 30% sites that are shared might be due to the retention of two KLF4 binding elements (at P19 and P40) on the wtAAV2^ΔKLF4^ virus. The statistical significance of this intersection was computed and presented in Fig. 4D.

**Figure 4.**
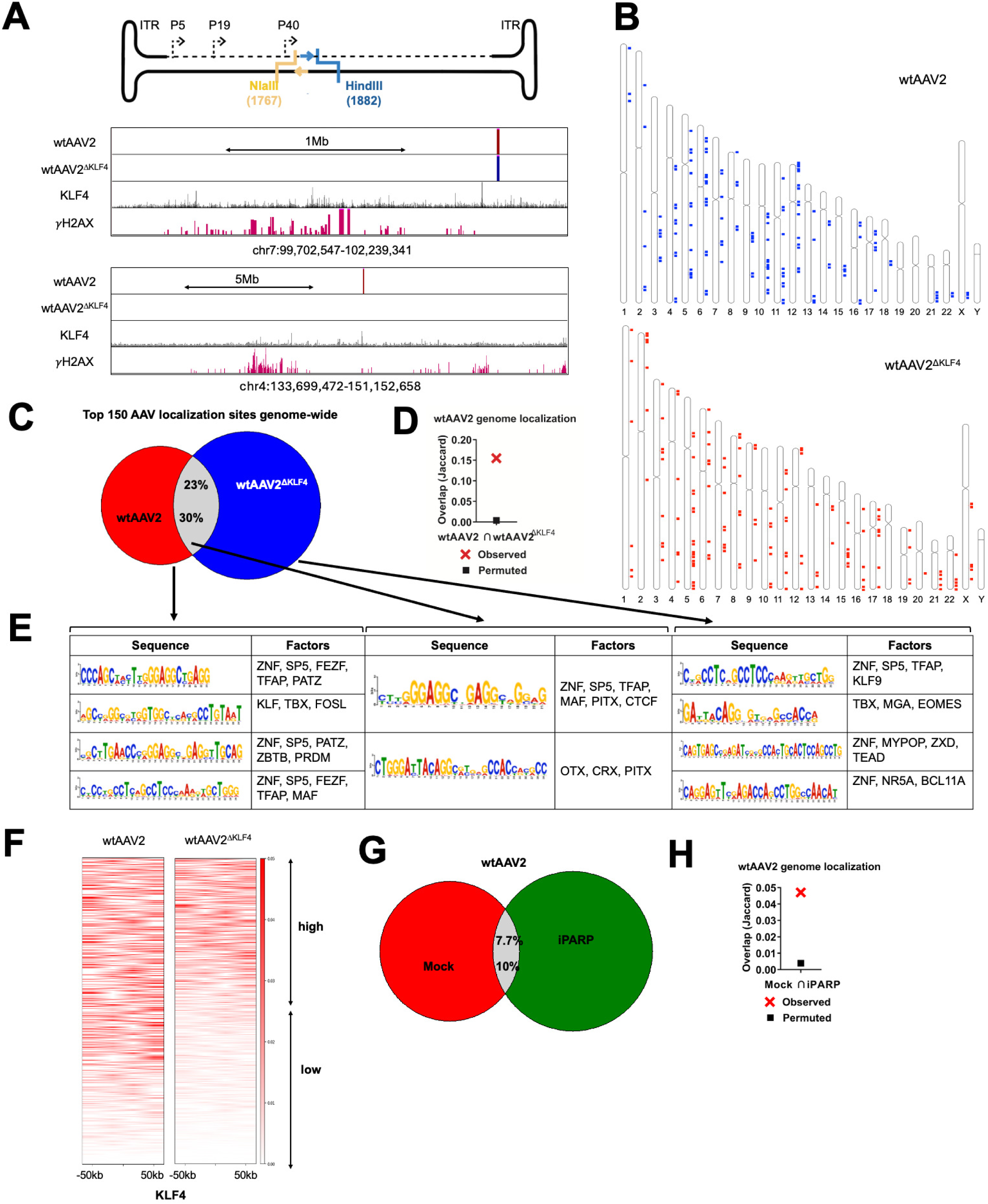
KLF4 binding site deletion attenuates wtAAV2 localization to DDR sites genome-wide. (A) (*top*) Schematic of wtAAV2 genome with the location of the HindIII site (blue) and NlaIII site (yellow) used for inverse PCRs indicated by forward (blue arrow) and reverse (yellow arrow) primers respectively. (*middle and bottom*) Representative UCSC genome browser tracks on human Chromosome 7 and Chromosome 4 of the indicated wtAAV2 variants identified using V3C-seq and compared with previously published KLF4 ChIP-seq in MCF7 cells^46^ and γH2AX in 293T cells^16^. (B) Genome-wide localization of the wtAAV2 (top) and wtAAV2^ΔKLF4^ (bottom) focused on the top 150 virus-associated HindIII fragments on the human genome. (C) Intersection of the genome coverage of the top 150 HindIII-localization sites of wtAAV2/wtAAV2^ΔKLF4^ viruses, with the (D) statistical significance depicted by Jaccard analysis. Jaccard values represent the extent of overlap, with 0 indicating no overlap and 1 indicating complete overlap. The “permuted” samples are computed by intersecting the V3C-seq data with a randomly generated library of 150 fragments of 5 kb size across the human genome. (E) Results of the *in-silico* analysis using MEME and TOMTOM^80^ pipelines of the fragments that are represented in the distinct subsets in Fig. 4C. The consensus sequences are presented in the left column and the corresponding transcription factors that have binding sites in the respective motifs are presented on the right columns. (F) Genome-wide profiling of wtAAV2 (left) and wtAAV2^ΔKLF4^ (right) localization sites relative to all KLF4 binding sites identified by ChIP-seq within 100 kb bins. These heatmaps were computed using the deeptools resource on the Galaxy project server^84^. (G) Venn diagrams showing the extent of overlap of the top 150 wtAAV2 localization sites on the human genome in the absence and presence of the PARP inhibitor Olaparib (labelled as iPARP). (H) Statistical analysis of the overlap was performed using Jaccard analysis (BEDtools suite^79^) using the same principle as described above.

To determine which host factors are enriched at the wtAAV2/wtAAV2^ΔKLF4^-associated sites, we performed *in-silico* analysis of the subsets of host genomic regions represented in Fig. 4C (red, grey and blue sections of the Venn diagram respectively). As shown in Fig. 4E (left-most columns), the consensus sequences that were enriched in wtAAV2-associated genomic regions (colored red in the Venn diagram) contained binding elements for transcription factors such as the ZNF family, SP5, KLF family and TFAP. Among these protein families, the ZNF, KLF and TFAP proteins are associated with cellular DNA damage and host genome instability^51–53^. The regions that were shared between wtAAV2 and wtAAV2^ΔKLF4^ (grey in the Venn diagram) were additionally enriched in binding elements for CTCF (Fig. 4E, middle columns), which is a host architectural protein that also associates with the cellular response to DNA damage^54, 55^. These shared genomic regions were also enriched in binding elements for TRX and PITX, which are involved in stress-induced responses^56, 57^. The cellular sites that were strongly associated exclusively with the wtAAV2^ΔKLF4^ virus (blue region of the Venn diagram) were also enriched in binding elements for ZNF, SP5 and TFAP proteins (Fig. 4E, right columns). This suggested that wtAAV2 viruses likely deploy redundant pathways to associate with cellular genomic regions. Strikingly however, these consensus sequences also contained binding elements for KLF9, but no other KLF family members. These observations suggested that in the absence of KLF4 binding to the 5’ end of the viral genome, wtAAV2^ΔKLF4^ compensates by associating with a different KLF family member.

Since KLF4-mediated nuclear foci bring together distal elements of the host genome^47, 58^, we compared all genome-wide localization sites of wtAAV2/wtAAV2^ΔKLF4^ to those of KLF4 ChIP-seq peaks^16, 59^. The localization of wtAAV2^ΔKLF4^ was attenuated in 50% of these host genomic regions (Fig. 4F, see high versus low peaks). In combination with the laser micro-irradiation studies described above, these observations suggested that there is a connection between KLF4 binding and wtAAV2’s ability to navigate the nuclear compartment.

KLF4 is known to associate with and be modified by the cellular DDR factor PARP1^36, 37^. We therefore assessed the impact of wtAAV2-localization to host genomic sites upon being treated with the PARP-inhibitor Olaparib (labelled as iPARP)^60^. As shown in Fig. 4G, only 10% of wtAAV2-associated genomic regions were shared with those of wtAAV2-infected cells treated with iPARP. Fig. 4H demonstrates the statistical significance of this intersection. Taken together, our findings showed that loss of KLF4 binding attenuates the ability of wtAAV2^ΔKLF4^ to localize efficiently to host DNA break sites genome-wide, likely through signals that are regulated by PARP.

### wtAAV2-induced PARP1 activity is necessary for viral gene expression and localization to cellular DDR sites

KLF4 has previously been shown to associate with PARP and PARylated cellular substrates at host telomeres^36^. Infection of 293T cells with wtAAV2 genomes at 5,000 vg/cell for 24 hours led to the PARylation of cellular substrates (shown by the marked smears in Fig. 5A, column 2 versus 1). Treatment of wtAAV2-infected cells with the pan-PARP1/2 inhibitor Olaparib (iPARP) led to a global attenuation of PARylated substrates (Fig. 5A, column 3 quantified relative to tubulin on the right). Moreover, the presence of iPARP in wtAAV2-infected 293T cells decreased *Rep 68/78* gene expression (Fig. 5B). This treatment also significantly reduced both KLF4 and PARP1 binding to the wtAAV2 genome in ChIP-qPCR assays (Fig. 5C, left and right respectively). iPARP also attenuated the laser micro-irradiation-induced relocalization of wtAAV2 genomes to DDR sites (Fig. 5D, 5E). Together with PARP1 ChIP-qPCR data showing a significant decrease in PARP1 association with the viral genome in the absence of the KLF4 binding element (Fig. 3D), these observations suggested that KLF4 binding sites drove the recruitment of PARP1 to wtAAV2. To dissect the role of PARP1 in regulating wtAAV2 genome localization and viral gene expression, we generated PARP1-deficient U2OS cells using CRISPR/Cas9-mediated genome editing (Fig. 5F). These PARP1-deficient cells were transiently transfected with either wild-type PARP1, PARP1^H909A^ and PARP1^T824A^ (deficient in KLF4 binding) or PARP1^E988K^ (deficient in catalytic activity). These complemented cells were then infected with wtAAV2 to examine PARP1’s ability to rescue wtAAV2 gene expression. *Rep68/78* gene expression was negligible in PARP1^-/-^ U2OS cells infected with wtAAV2 virus (Fig. 5G, pUC18 transfected column). Complementing wild-type PARP1 activity rescued wtAAV2 gene expression in these cells (Fig. 5G, PARP1 column). However, overexpression of PARP1 mutants unable to interact with KLF4 (PARP1^H909A^, PARP1^T824A^) did not rescue *Rep68/78* gene expression. Surprisingly, the PARP1 mutant lacking catalytic activity (PARP1^E988K^)^36, 37^ showed a partial rescue of wtAAV2 gene expression (Fig. 5G, right column). We confirmed that all the transfected plasmids expressed stable PARP1 proteins (Fig. 5H). Taken together, these studies demonstrated that the interaction between KLF4 and PARP1, and likely not PARP1’s catalytic role in PARylation, that is necessary for efficient wtAAV2 gene expression.

**Figure 5.**
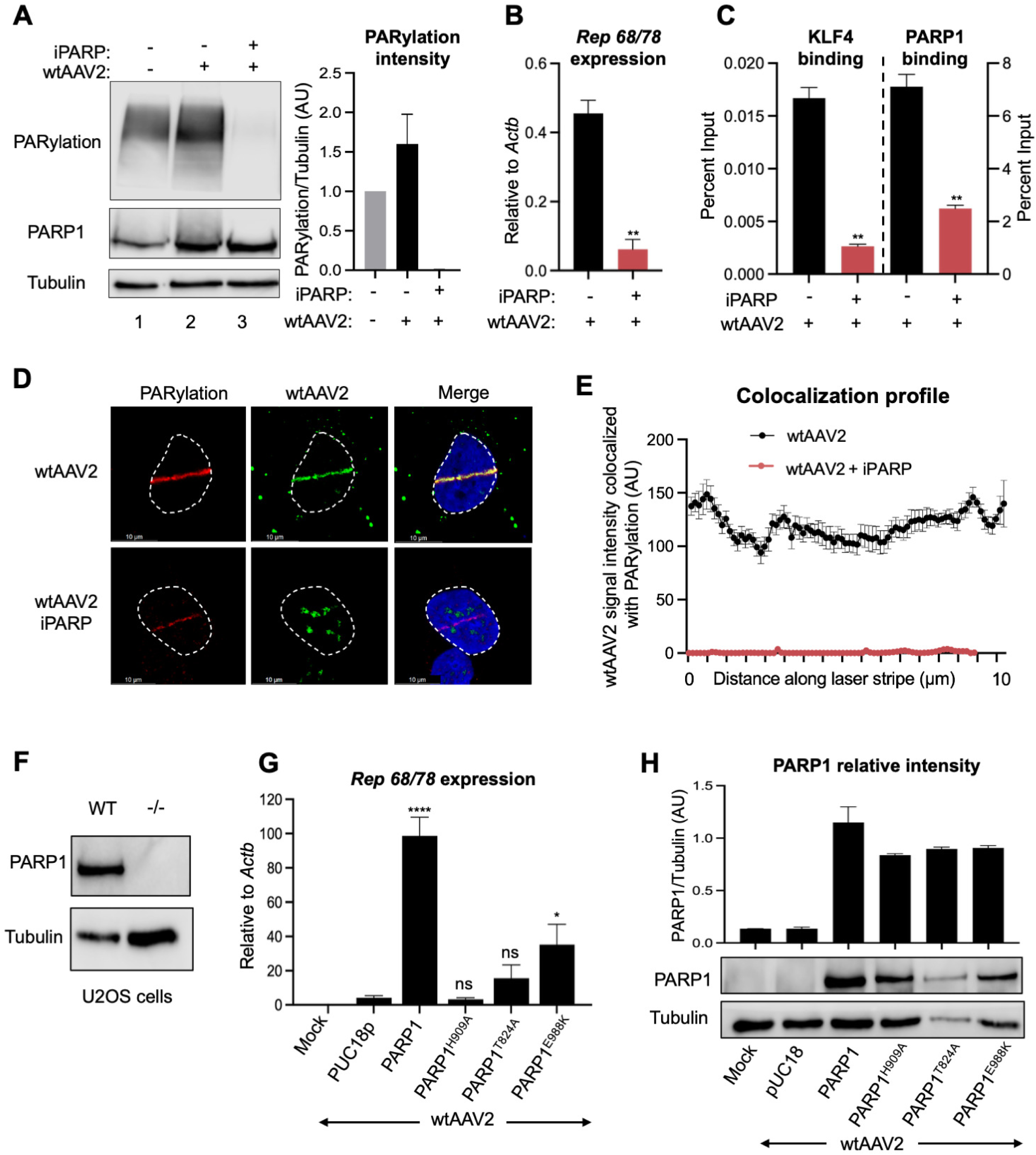
wtAAV2-induced PARP1 activity is necessary for viral gene expression and localization to cellular DDR sites. (A) Western blot analysis of the impact of wtAAV2 infection of 293T cells at an MOI of 5,000 vg/cell at 24 hpi. The PARP1 and PARylation levels were measured with the respective antibodies and Tubulin was used as loading control. The figure on the left is a representative immunoblot and the right shows the average of PARylation levels relative to Tubulin in 3 independent biological replicates of viral infection. Error bars represent SEM of PARylation to Tubulin ratios. (B) RT-qPCR analysis of *Rep 68/78* transcript levels in wtAAV2 infected 293T cells in the presence of Olaparib (PARPi). The corresponding impact on (C) KLF4 binding (left) and PARP1 binding (right) at P5 was measured using ChIP-qPCR analysis. Statistical analysis was performed with a t test, ** representing p < 0.01. (D) Impact of Olaparib treatment (PARPi) on wtAAV2 localization to the cellular sites of DNA damage was evaluated using laser micro-irradiation coupled with Immuno-FISH in wtAAV2 infected U2OS cells at 24 hpi. The DNA damage site was monitored by PARylation staining (red) and the viral genomes were monitored by fluorophore-labelled oligos (green). The nucleus was monitored by DAPI staining (blue) and the nuclear borders are demarcated by dashed white lines. (E) The profile of wtAAV2 colocalizing with PARylation signal across multiple micro-irradiated nuclei was plotted in 75 discrete intervals along the DDR site. Data is presented as mean ± SEM of the colocalized intensity in at least 10 nuclei. (F) PARP1 western blot in wild-type and PARP1-deficient U2OS cells demonstrating knockout of the PARP1 protein with Tubulin as loading control. (G) RT-qPCR of *Rep68/78* transcripts in PARP1-deficient U2OS cells that were complemented with the indicated PARP1 expression vectors for 24 hours before being infected with wtAAV2 for 24 hpi at an MOI of 5,000 vg/cell. Data is presented as mean ± SEM of expression relative to *Actb* in three independent replicates, with statistical significance computed using t tests, * representing p < 0.05, **** representing p < 0.0001 and ns denoting no statistical significance. (H) The protein stability of PARP1 during ectopic expression of PARP1-deficient cells was monitored by western blots 24 hours after transient transfection with the expression vector, and the mean ± SEM of three independent replicates of PARP1 to Tubulin ratios are presented on the top. Representative western blots are shown in the bottom.

### KLF4 binding element is sufficient to increase rAAV2 transgene expression and colocalization with cellular DDR foci

To determine whether KLF4 binding elements are sufficient to increase the expression of rAAV2 vectors, we inserted a dimer of the KLF4 consensus binding sequence (AGGGGTGGAGTC) between the ITR and the CBE enhancer in an rAAV2 vector expressing a GFP reporter transgene (Fig. 6A). Co-transfection of these prAAV2 plasmids into 293T cells with pHelper and pRepCap2-expressing plasmids revealed that rAAV2^KLF4^ genomes were not produced more efficiently than rAAV2 (Fig. S1). When transgene expression was monitored by FACS analysis, these rAAV2^KLF4^ vectors expressed GFP four-fold more efficiently in 293T cells (shown in representative FACS plots in Fig. 6B). This increased expression was reproducible when monitoring the mean fluorescent intensity (Fig. 6C) or percent GFP positive cells (Fig. 6D) at 24hpi. When immunolabelled in rAAV2^KLF4^-transduced U2OS cells, PARylated proteins colocalized with the fluorophore-conjugated vector genomes in small foci in the nuclear compartment (Fig. 6D, shown by white arrows). Taken together, our findings suggest that KLF4 binding is sufficient to facilitate the colocalization of rAAV2 genomes with cellular DDR sites, thereby facilitating efficient transgene expression.

**Figure 6.**
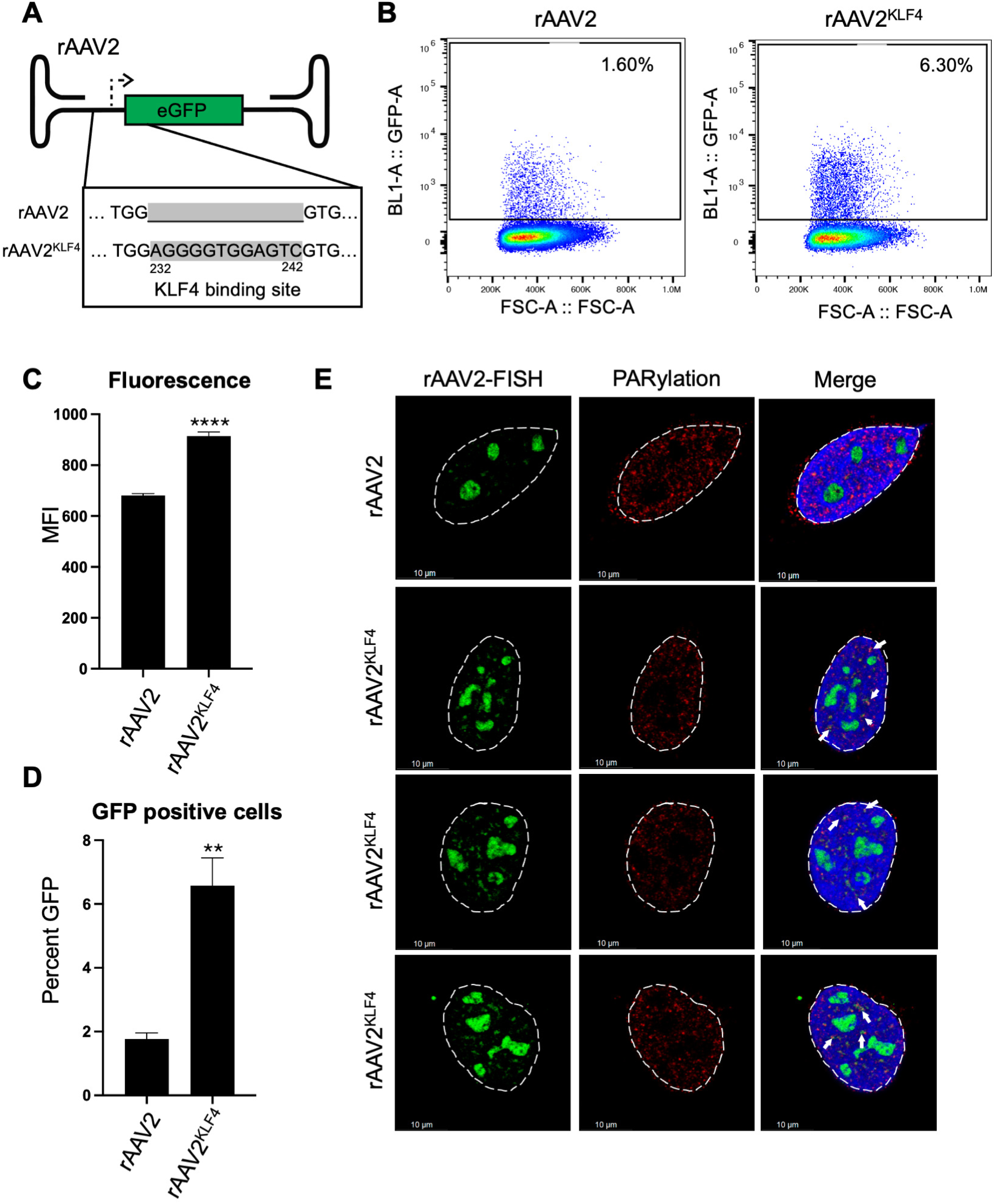
KLF4 binding element is sufficient to increase rAAV2 transgene expression and colocalization with cellular DDR foci. (A) Schematic of the rAAV genomes showing the location of the insertion of the KLF4 binding element dimer sequences (represented by grey highlights). (B) Impact of rAAV2/rAAV2^KLF4^ transduction on 293T cells transduced at an MOI of 5,000 vg/cell at 24 hpi. The figure demonstrates representative FACs plots from 3 independent experiments in live cells identified by forward and side-scatter analysis followed by assessment for GFP positivity. The percent positive cells are shown in the inset. (C) Average mean fluorescence intensity of GFP expression in 293T cells transduced with the indicated rAAV vectors for 24 hpi at an MOI of 5,000 vg/cell. (D) These cell populations were assessed for GFP positivity in live cells using FACS analysis. Data is presented as mean ± SEM of cells from at least three independent transduction experiments with statistical analysis performed using t tests. P values are computed as: ** representing p < 0.01 and **** representing p < 0.0001. (E) Representative Immuno-FISH images of U2OS cells transduced with the indicated rAAV vectors at an MOI of 5,000 vg/cell for 24 hpi followed by co-staining for vector genomes (green), PARylation (red) and DAPI (blue) to demarcate the nuclei. The white dashed lines indicate the nuclear borders and the white line represents 10 micrometers. White arrows in the nuclei indicate regions of colocalization between the vector genome and PARylation.

## DISCUSSION

wtAAV2 exploits host DNA damage response pathways to localize its genome to nuclear compartments that support persistence and viral gene expression. In this study, we demonstrate that the transcription factor KLF4 and the DNA damage response (DDR) protein PARP1 cooperatively regulate wtAAV2 genome localization during non-productive infection. Many wtAAV2-associated host loci are proximal to nucleolar chromatin, where rDNA genes cluster. KLF4 enrichment at these rDNA regulatory regions provides a plausible bridge between KLF4-bound chromatin and DDR-marked sites. Disruption of KLF4, by siRNA knockdown or targeted deletion of a KLF4-binding element on the viral genome, reduces *Rep68/78* gene expression and diminishes recruitment of wtAAV2 genomes to laser micro-irradiated DDR sites. Furthermore, inhibition of PARP activity with Olaparib (iPARP) reduces detectable KLF4 and PARP1 occupancy on the wtAAV2 genome and alters its nuclear localization pattern. These findings suggest that the association of wtAAV2 genomes with distinct nuclear compartments is closely linked to PARP1-mediated modification of the local host factors that aid the viral life cycle. Together, our findings support a model in which KLF4–PARP1 interactions position wtAAV2 genomes at DDR-enriched chromatin sites that are permissive for viral transcription.

Our findings build on previously established roles of KLF4 in regulating the life cycle of the DNA tumor viruses EBV and HPV. KLF4 regulates the differentiation-dependent lytic cycle of EBV by regulating the activity of the promoters of BZLF1 and BRLF1 genes^39^. In HPV31, KLF4 binds the upstream regulatory region (URR) and cooperates with BLIMP1 to control viral transcription^38^. Interestingly, in both systems, KLF4 also functions as a key cell cycle regulator that controls the differentiation-dependent amplification of host cells. However, in our system of wtAAV2 monoinfection of immortalized cell lines (U2OS and 293T), KLF4 is not required to maintain the differentiation status of the host cells. In the absence of these functions, the ability of KLF4 to interact with PARP1 seemingly enables the formation of nuclear viral reservoirs. Strikingly, this function of transporting the viral genome to cellular sites of DNA damage has previously been reported to be performed by NS1, the essential phosphoprotein that is required for the lifecycle of the autonomous parvovirus MVM^24^. Since NS1 and the REP 68/78 proteins are structurally and functionally similar, it remains unknown why the ability to transport the viral genome to cellular DDR sites is not within the purview of the REP 68/78 protein. One possibility is that wtAAV2 infection does not generate high levels of REP 68/78 proteins in the absence of a helper virus, which might limit their localization function. Alternatively, this might be a redundant mechanism that wtAAV2 has evolved to usurp another host factor for essential viral functions. Lastly, it is noteworthy that just like KLF4-drives wtAAV2 localization to cellular DDR sites, its cooperative Yamanaka factor OCT4 has been demonstrated to associate with HPV-E7 to regulate the formation of cervical cancers^61^. These phenomenon in distinct DNA viruses might reflect their convergent evolution in usurping essential transcription factors for viral success.

KLF4’s known ability to form nuclear biomolecular condensates (BMCs) is hypothesized to be a driver of nuclear reorganization that brings together distal elements on the eukaryotic genome^47, 58^. Consistent with this model of KLF4 activity, global knockdowns led to an attenuated localization to induced cellular DDR sites. Strikingly however, in the wtAAV2^ΔKLF4^ genome, localization was partially inhibited, possibly due to the retention of the KLF4 binding elements in P19 and P40 promoter regions. Of note, condensate formation by wtAAV2 has only been observed under replicative conditions and involves accessory proteins such as MAAP^62, 63^, which are absent in our non-replicative system. An alternative possibility is that KLF4-mediated chromatin remodeling (potentially driven by the ability of KLF4 to interact with epigenetic modifiers like P300^64, 65^) draws wtAAV2 to DDR-enriched microenvironments, whereas autonomous parvoviruses such as MVM encode NS1 to perform these targeting functions directly. Discriminating between condensate-dependent recruitment and classic protein–DNA tethering will require perturbations that disrupt phase-separation (e.g., 1,6-hexanediol treatment), live-cell imaging of KLF4/PARP1 localization and FRAP-based assays. Some of these properties of DNA viruses to generate nuclear condensates have recently been demonstrated for Adenoviruses^66^.

The over-representation of KLF4 binding elements at cellular sites of DNA damage raises the question of whether KLF4 plays a role in the initial induction of cellular DNA breaks. One plausible route is by the generation of transcription–replication conflicts. KLF4-dependent chromatin remodeling may elevate local transcription in ways that intersect with replication stress, indirectly generating DNA damage. Indeed, replication stress sites, generated during wtAAV2 infection^17^, might be the initial events that activate PARP1 leading to PARylation of local substrates. Consistent with this prediction, we observe increased cellular PARylation during wtAAV2 infection and reduced PARP1 engagement with the viral genome upon Olaparib treatment. Because Olaparib treatment and inoculation were simultaneous, PARP1 may become sequestered on host chromatin by the inhibitor, which could account for reduced PARP1 on the viral genome. Future work should map PARylated substrates and their genomic distribution during wtAAV2 infection to define whether PARP1 activity precedes or follows KLF4-mediated recruitment. Our findings build upon prior mass spectrometry studies that had discovered PARP1 as one of the binding partners of REP 68/78 during wtAAV2 infection^40^. We propose that PARP1, through its interaction with KLF4, is specifically required for wtAAV2 gene expression. These observations resemble EBV, where PARP1 activity restricts lytic cycle by supporting latent viral gene expression^31–33^. Similarly, PARP1-mediated inhibition of cellular DDR signals is deployed by Adenovirus for E4orf4-induced cancer cell death^67^. On the other hand, PARP1 activity is known to facilitate the degradation of cellular Interferon Alpha Receptor proteins, facilitating efficient Influenza A virus replication^68, 69^. These activities might point towards distinct protein degradation functions of PARylation, which are yet to be deciphered for wtAAV2 infection.

The ability of KLF4 and PARP1 to enhance rAAV gene therapy vector expression raises exciting possibilities about how they regulate the fate of wtAAV2 virus genomes and rAAV vectors. By regulating the positioning of the virus/vector genomes within the nuclear microenvironment, they can actively regulate their ability to express (as shown in this study). Of note, the KLF4 binding element is directly adjacent to the D element on the wtAAV2 genome, which is known to enhance rAAV2 packaging and vector transduction efficiency^70–72^. Future investigations could investigate how these interactions regulate the ability of wtAAV2/rAAV to form extrachromosomal concatemers^73^ that serve as templates for long-term viral gene expression, vector persistence, and impact on integration^74^. Importantly, our studies focus on interrogating the fate of wtAAV2/rAAV2 genomes during monoinfection, which differs from replicative infection in the presence of helper viruses that cause large-scale nuclear reorganization. Therefore, our observations are more closely representative of the impact of gene therapy vectors on transduced target cells. Taken together, our findings provide broad implications for virology, gene therapy, and host-pathogen interactions by leveraging the ability of rAAV2 to mimic wtAAV2’s mechanisms to access cellular DDR sites.

## MATERIALS AND METHODS

### Cell lines

Female Human Embryonic Kidney (HEK293T) cells and female human U2OS osteosarcoma cells were maintained in Dulbecco’s Modified Eagle’s Medium [DMEM, high glucose (Gibco)] supplemented with 5% Serum Plus (Sigma-Aldrich) and 50 µg/mL gentamicin (Gibco). The cells were cultured at 37 °C in 5% CO₂ incubators. Cells were routinely tested for mycoplasma contamination using the MycoStrip Mycoplasma Detection Kit (InvivoGen) and the basal DNA damage levels monitored by γH2AX staining.

### Virus and vector preparation

The plasmid infectious clones of wild-type AAV2 (wtAAV2) and AAV2^ΔKLF4^ (wtAAV2 ^ΔKLF4^) were transfected into producer HEK293T cells using Polyethylenimine (PEI; Polysciences). The plasmid infectious clones (derived from SSV9) were co-transfected in HEK293T producer cells by triple or double transfection with pHelper plasmids (expressing Adenovirus E2A, E4 and VA-RNA) and pRepCap2 (in the case of rAAV vector production). These transfections were performed at molar ratios of 1:1:1 (pHelper: prAAV: pRep/Cap2) or 1:1 (pHelper: pwtAAV2) for a total of 30µg of DNA with 3µl of PEI for each microliter of DNA. Virus was collected 60 hours post-transfection and cells were lysed by seven freeze-thaw cycles in liquid nitrogen. The clarified lysate was treated with DNase I (Thermo Scientific) to degrade unencapsidated DNA. wtAAV2/rAAV2 genome titers were quantified by qPCR using primers complementary to the respective ORFs using serial dilutions of the plasmid genomes as standards. The virus (or vector) genome concentration was calculated as viral genomes per milliliter (vg/mL).

rAAV2 and rAAV2^KLF4^ vectors (containing a dimer of the KLF4 consensus sequence between the ITR and CBE enhancer) were produced analogously and genome integrity of the plasmids were first confirmed by agarose gel electrophoresis, restriction enzyme digest and Sanger sequencing across the insert.

### RNAi, CRISPR guide RNA and Inhibitor treatment

For KLF4 knockdown, cells were transfected with two independent siRNAs targeting KLF4 or a non-targeting siMock control using Lipofectamine RNAiMAX transfection reagent (Thermo Scientific) at a final siRNA concentration of 10nM. Knockdown was carried out for 24 hours prior to infection. The KLF4 siRNAs used were s17793 and s17794 from Thermo Scientific.

PARP1-deficient U2OS cells were generated by co-transfecting equimolar amounts of 3 guide RNAs targeting the PARP1 locus using Lipofectamine CRISPRMAX Cas9 Transfection Reagent (Thermo Scientific). At 72 hours post-transfection, the bulk population of transfectants were single-cell cloned into 96-well plates using limiting dilution cultures. The individual clones were evaluated for PARP1 protein knockout at 15 days post-dilution with Tubulin as loading control.

PARP inhibition was carried out using the pan-PARP1/2 inhibitor Olaparib (AZD2281; Selleck Chemicals S1060), applied at a concentration of 1 µM concurrently with virus inoculum at the time of infection and maintained throughout the time-course of infection. Vehicle controls received matched DMSO, which is the medium for dissolving the Olaparib chemical. Where indicated, alternative PARP1 constructs (WT and mutants defective for catalysis or KLF4 interaction) were transiently expressed 24 h pre-infection. Mutations were made on the expression plasmid backbones pEGFP-N1-PARP1 (H909A/T824A/E988K) that are described below and in the primer table.

### Antibodies

Primary antibodies used in this study and their respective dilutions used in western blots were: γH2AX (Abcam, ab11174, 1:1000); α-Tubulin (Millipore, 05-829, 1:5000); REP68/78 (IF11.8, 1:1000); PARP1 (BD Biosciences, 556494, 1:1000); KLF4 (Cell Signaling Technology, 4038S, 1:1000); PARylation (Cell Signaling Technology, 89190, 1:1000).

Secondary antibodies used for imaging western blots in this study and their respective dilutions were: HRP-conjugated anti-mouse (Cell Signaling, 7076S, 1:5000) and anti-rabbit (Cell Signaling, 7074, 1:5000).

### Plasmids

The PARP1 expression vector (pEGFP-N1-PARP1; Addgene plasmid # 211578) was obtained from Addgene and was a gift from Dr. Chris Lord^75^. Site directed mutagenesis was performed on the PARP1 open reading frame using the primers indicated in the primer table.

### Western Blot

Cells were seeded at a density of 5 × 10⁵ cells per well in 6-well plates and infected with wtAAV2 or wtAAV2^ΔKLF4^ mutant viruses at a multiplicity of infection (MOI) of 5,000 viral genomes per cell. After 24 hours post-infection, cells were washed with cold PBS and lysed on ice for 15 minutes using RIPA buffer (50 mM Tris-HCl pH 7.4, 150 mM NaCl, 1% NP-40, 0.5% sodium deoxycholate, 0.1% SDS) supplemented with protease inhibitors (MCE), 1 mM sodium orthovanadate, and 10 mM sodium fluoride. Lysates were clarified by centrifugation at 13,000 rpm for 10 minutes at 4°C. Protein concentration was determined using the BCA Protein Assay Kit (Bio-Rad). Equal amounts of protein (typically 30–40 µg) were mixed with 6X Dye Loading, boiled at 95°C for 5 minutes, and loaded on 10% SDS-PAGE gels. Proteins were transferred to PVDF membranes (Bio-Rad) using semi-dry transfer at 25 V for 30 minutes. Membranes were blocked in 5% non-fat dry milk in TBST (Tris-buffered saline with 0.1% Tween-20) for 30 minutes at room temperature, followed by incubation with primary antibodies overnight at 4°C.

After washing, membranes were incubated with HRP-conjugated secondary antibodies (1:5000) for 1 hour at room temperature. Signal detection was carried out using ECL substrate (Bio-Rad) and imaged using a Li-COR Odyssey system. Band intensities were quantified using ImageStudio software (Li-COR).

### Reverse Transcription combined with quantitative PCR (RT-qPCR)

Total RNA was extracted using PureZOL reagent (Bio-Rad) followed by chloroform-mediated phase separation and isopropanol precipitation. RNA samples were treated with RNase-free DNase I (Promega) at 37°C for 30 minutes prior to reverse transcription. Reverse transcription was performed using the iScript cDNA synthesis kit (Bio-Rad) with 1 µg of RNA in a 20 µL reaction. qPCR was conducted using SYBR Green Master Mix (Bio-Rad) on a CFX96 Real-Time PCR System under the following cycling conditions: 95°C for 5 min, followed by 40 cycles of 95°C for 10 sec and 60°C for 30 sec. Relative expression was calculated using the ΔCt method, normalized to *Actb* loading controls. No-RT controls were used to measure any residual viral DNA carry-over; and these signals were used to establish the baseline during RT-qPCR analysis. The formation of single amplicons were confirmed by melt-curve analysis of the qPCR assays.

### Chromatin Immunoprecipitation combined with quantitative PCR (ChIP-qPCR)

Cells were crosslinked with 1% formaldehyde for 10 minutes at room temperature and quenched with 0.125 M glycine. Cells were lysed in an SDS ChIP Lysis Buffer for 20 minutes on ice (1% SDS, 10 mM EDTA, 50 mM Tris-HCl, pH 8, and protease inhibitor) before the chromatin was sheared to ∼500 bp fragments using a Diagenode Bioruptor Pico (120 cycles of 30 sec on/off). The nucleo-protein complexes were incubated with antibodies immobilized on Protein A Dynabeads (Thermo Scientific) overnight at 4°C using 2 µg of the respective antibodies antibody (KLF4, PARP1). After the overnight incubation, beads were washed sequentially with low salt wash (0.01% SDS, 1% Triton X-100, 2 mM EDTA, 20 mM Tris-HCl pH8, 150 mM NaCl), high salt wash (0.01% SDS, 1% Triton X-100, 2 mM EDTA 20 mM Tris-HCl pH8, 500 mM NaCl), LiCl wash (0.25M LiCl, 1% NP40, 1% DOC, 1 mM EDTA, 10 mM Tris HCl pH8), and TE buffers. DNA was eluted in 1% SDS, 0.1 M NaHCO₃ and reverse crosslinked overnight at 56°C with 0.2 M NaCl and Proteinase K (NEB). DNA was purified using a Qiagen PCR Purification Kit and eluted in 100 µl of Buffer EB (Qiagen). The pulldown DNA was analyzed by qPCR using primers targeting viral and host loci across the wtAAV2 genome as indicated (Table 1).

### Immunofluorescence Imaging

wtAAV2-infected U2OS cells grown on 35 mm coverslips were harvested at 24 hpi by pre-extracting with CSK buffer (10 mM PIPES, pH 6.8, 100 sodium chloride, 300 mM sucrose, 1 mM EGTA and 1 mM magnesium chloride) for 3 minutes followed by CSK buffer containing 0.5% Triton X-100 for 3 minutes at room temperature. Cells were fixed with 4% paraformaldehyde (EMS) for 10 minutes, washed with PBS and permeabilized with 0.5% Triton X-100 for 20 minutes at room temperature. The samples were blocked with 3% BSA in PBS for 30 minutes, incubated with the indicated primary antibodies (γH2AX, PARylation, KLF4, PARP etc) for 30 minutes to 1 hour at room temperature, washed with PBS and incubated with Alexa Fluor-conjugated secondary antibodies (1:1000 dilution) for 30 minutes in the dark. The samples were washed in PBS and counterstained with DAPI-containing Flouromount (SouthernBiotech) on glass slides. Imaging was performed using a Leica Stellaris confocal microscope with a 63X oil immersion objective.

### Immunofluorescence coupled with Fluorescent In-Situ Hybridization (Immuno-FISH)

The Immuno-FISH assays were performed on U2OS cells that were plated and infected as described for the immunofluorescence assays above. DNA probes complementary to the wtAAV2 genome were purchased from IDT and were labeled using 250 mM Aminoallyl-dUTP (Thermo Scientific) and TdT (Promega). The labelling reactions were performed overnight at 37 degrees Celsius, inactivated at 70 degrees Celsius for 10 minutes and precipitated using isopropanol. The oligos were washed in 75% ethanol and dissolved in 15 µl of 0.1 M sodium bicarbonate (pH 8.3). 0.75 µl of 20mM NHS esters (Thermo Scientific) were conjugated to the labelled oligos for 2 hours in the dark, precipitated in isopropanol and cleaned up using a PCR clean-up column (Promega). Cells were pre-extracted with CSK buffer and CSK buffer containing 0.5% Triton X-100 for 3 minutes each at room temperature as described above before being fixed with 4% paraformaldehyde, and permeabilized. Samples were RNAse treated in 2X SSC for 1 hour at 37 degrees Celsius and denatured in 10% formamide solution in 2X SSC for 2 hours at 37 degrees Celsius. The samples were hybridized with 1 µl of labelled probe in 20 µl of hybridization buffer overnight at 37°C in a humidified chamber. Post-hybridization washes were performed in 2X SSC with 0.1% Triton X-100 at 37 degrees Celsius 3 times, 3 washes in 2X SSC and immunostaining was performed as described above. Samples were mounted with DAPI and imaged using a Leica Stellaris Confocal microscope equipped with a 63X oil objective lens.

### Laser Micro-Irradiation

Cells were seeded on glass-bottom dishes and processed as previously published^24, 25^. Five minutes prior to irradiation, cells were sensitized with 3 µL of Hoechst 33342 dye (Thermo Scientific), which was added directly to the culture medium. Micro-irradiation was carried out using a Leica Stellaris confocal microscope equipped with a 405 nm laser and a 63X oil immersion objective lens with 3X digital zoom. Laser settings were calibrated to 25% power at 40 Hz frequency, and irradiation was applied for 1 frame per field of view. Regions of interest (ROIs) were manually selected within the nuclear compartment, ensuring that laser tracks did not cross the nuclear envelope. Immediately following irradiation, cells were fixed and processed for immunofluorescence and immuno-FISH imaging as described in the respective sections above.

The colocalization of FISH probes with induced cellular DNA damage was quantified by measuring the intensity of FISH probe-associated signal along the laser stripe (monitored by PARylation or H2AX staining). The FISH signal intensity along the induced DNA break was quantified using the plot profile tool on the FIJI software. Signal intensities were measured at defined intervals along the region of interest (ROI) of the irradiated nuclei. These values were measured for 20-40 nuclei in 3 biological replicates of viral infection of U2OS cells.

### Viral Chromosome Conformation Capture coupled with Sequencing (V3C-seq)

V3C-seq was performed as previously performed for wtAAV2^16, 76^ with slight modifications. Briefly, 10 million wtAAV2 infected cells at 24 hpi were crosslinked with 1% formaldehyde for 10 minutes, quenched with glycine, and lysed. Chromatin was digested with the primary restriction enzymes (HindIII; NEB) and ligated under dilute conditions to favor intramolecular ligation. DNA was purified, secondary digested using NlaIII (NEB), circularized and purified. The interactome of the wtAAV2 genome was measured using inverse PCR primers and nested inverse PCR primers complementary to the wtAAV2 genome. Libraries were prepared from 1 microgram of DNA using NEBNext Ultra II DNA Library Prep Kit and sequenced using single-end sequencing on an Illumina NextSeq platform spiked with 25% phiX spike-in to increase library complexity.

High-throughput sequencing data was aligned to the hg38 human reference genome using the Minimap2 alignment program^77^. The aligned data was sorted using Samtools^78^ and genome-wide coverage was computed using BEDtools^79^. Independent biological replicates of the V3C-seq studies were intersected using BEDtools. The aligned reads were computed against a library of all HindIII fragments on the hg38 genome using RStudio. The top 150 localization peaks were used to perform comparative intersections using BEDtools. The motifs that are enriched at the shared and unique localization sites were computed using the MEME motif search platform^80^. The transcription factors associated with the most highly-represented motifs were measured using the FIMO and TOMTOM^81^ platforms. Statistical analysis of the intersection was performed using Jaccard analysis on BEDtools. The “observed” values were calculated using intersection of the wtAAV2 localization with that of wtAAV2^ΔKLF4^ localization sites. This intersection was computed relative to “permuted” values, where the wtAAV2 localization sites were intersected with a randomly generated set of genomic coordinates of the same average size and number as wtAAV2-associated regions. These computations were performed using the Galaxy project server^82^.

### FACS analysis

Cells were transduced with GFP containing rAAV2 or rAAV2^KLF4^ at MOI 5,000 vg/cell. At 24 hpi, GFP positivity and mean fluorescence intensity (MFI) were assessed by flow cytometry on a BD LSR Fortessa using a 488 nm, blue laser. Forward-scatter and side-scatter was used to gate on lice cells and thereafter, the GFP-positive cells were assessed according to the presence of singlets. The computation of GFP-positive cells were gated and analyzed using FlowJo software (version 10.10).

### Statistical analysis and replicates

Unless otherwise indicated, all experiments were performed in at least three independent biological replicates of viral infection or vector transductions. Statistical analysis was performed using GraphPad Prism and the respective details of the statistical analysis are provided in the corresponding figure legends.

### Data Accession

All high-throughput sequencing data, including raw files and analyzed files, have been uploaded to the data repository at the NCBI Gene Expression Omnibus (GEO) and are accessible under the accession number GSE320210.

## ACKNOWLEDGEMENTS

This research was funded partially by NIH/NIAID K99/R00 Pathway to Independence Award, grant number AI148511, to K.M.; the Wisconsin Partnership Program’s New Investigator Award (PERC Grant G-4942) to K.M. and NIH/NIGMS R35 Maximizing Investigator’s Research Award (MIRA), grant number GM154938, to K.M. C.I.S.L. is funded by NSF Graduate Research Fellowship Program award DGE-2137424. R.R.A. was funded by a SciMED Graduate Research Scholarship from the University of Wisconsin-Madison. We thank the Precision Medicine Research Service of the UW Center for Human Genomics and Precision Medicine for high-throughput sequencing.

---

**Supplemental Figure S1: KLF4 element insertion into rAAV2 genomes do not impact vector replication.** Genome copies of rAAV2/rAAV2^KLF4^ vector genome monitored by qPCR analysis of prAAV/prAAV2^KLF4^, pHelper and pRepCap2 co-transfected 293T cells. Data is presented as median of three independent experiments monitoring vector genome production.

